# Core-Shell Hydrogel System to Protect the Enzyme Activity of Phytase from Environmental Stress

**DOI:** 10.1101/2025.03.07.642130

**Authors:** Eunhye Yang, Waritsara Khongkomolsakul, Younas Dadmohammadi, Alireza Abbaspourrad

## Abstract

Fortification of phytase from *Aspergillus niger* (phyA) in a vegetarian diet is a practical strategy to solve mineral deficiencies induced by the presence of phytate. To protect phyA’s activity and retention from environmental stress such as heat and acidic conditions, we evaluated the use of a core-shell hydrogel bead, where the core is composed of a phyA-chitosan complex, and the shell is formed by cross-linking alginate with κ-carrageenan. The phyA loading capacity is 52.2%, with high encapsulation efficiency (82.6%). When forming the hydrogel beads, a needle diameter of 0.5 mm can create a 2.5 mm bead. The beads were found to remain intact during dehydration under a vacuum at 30 ℃. The formed hydrogel beads protected 79.7% of the phyA activity after heating at 100 °C for 12 min. The beads protected their cargo against salt, pH changes, and protease. These results suggest that core-shell beads are suitable for protecting enzyme activity against various processing stresses, which makes them useful as a delivery method for phytase in food applications.

## 1. Introduction

Plant-based foods include a high level of phytate, which can chelate minerals in food, keeping them from being absorbed and, consequently, causing nutritional deficiencies. As a solution to mineral deficiency, phytase is a naturally occurring enzyme in the digestional tract that hydrolyzes phytate. However, the quantity of phytase in the human body is insufficient to degrade phytates before mineral absorption (Brouns, 2022), making it important for those following a plant-based diet to add additional phytase to their diet.

There are three challenges to using dietary phytase in food fortification. First, phytase should have high thermal stability to maintain enzymatic activity in practical applications, such as high cooking temperatures associated with maize porridge. Second, phytase should also have a high tolerance to various food components, including salt, pH, and other ingredients, such as glucose, which can bind with phytate. Third, to maintain enzyme activity in the human body, the phytase needs to maintain high activity in low pH levels (around pH 2) to be able to survive in the stomach.

In our previous study, we designed a core-shell hydrogel bead with heat resistance for enhanced thermal stability of phytase from wheat (E. Yang et al., 2024). We found that the thermal stability of phytase was improved when the core-shell structures were made of heat-resistant polysaccharides such as chitosan, alginate, and κ-carrageenan and that the system protected phytase activity (E. Yang et al., 2024). Because phytase from wheat has only one optimum pH and rapidly loses activity at pH 2, we searched other types and sources for phytase.

Compared to 6-phytase found in wheat, the type of phytase from *Aspergillus niger* (phyA) is 3-phytase. 3-phytase and 6-phytase are both enzymes that break down phytic acid, but they differ in where they initially attack the molecule; 3-Phytase hydrolyzes the phosphate group at the C3 position of phytic acid first (6-Phytase: C6 position). The initial hydrolysis site affects the enzyme’s interaction with the substrate and the stability of the intermediate products, influencing their efficiency under different pH conditions. As a result, phyA is active between pH 2 and 6.5 (6-phytase: pH 4-5.5) and thus has a higher stability against acidic conditions. 3-Phytases are typically more active in acidic conditions (pH ∼2) because their catalytic mechanism relies on the protonation of specific residues, facilitating the hydrolysis of the phosphate ester bond. Their secondary pH optimum at ∼6.5 may reflect adaptability to less acidic environments, where specific ionizable residues remain functional. This property makes it easier to use phyA in food applications, so we selected phyA instead of 6-phytase for further study.

The food matrix has many variables, such as temperature, salt, and pH, which can affect the enzyme activity and the structure of the encapsulation system (Saqib et al., 2022). For example, maize porridge, a representative food matrix of plant-based diets, generally includes 0.2 w/v% salt and a pH of 6.0 with a heating process over 10 minutes at 100 °C (Wang et al., 2024). Salt and pH significantly affect the swelling behavior of hydrogel beads and release properties of the cargo materials (Y. Wang et al., 2024). Also, heat treatment can cause the membrane of normal hydrogel beads to break down and eventually damage the encapsulated phyA.

We hypothesized that core-shell hydrogel beads would have a protective effect, improve thermal stability, and enhance phyA’s salt and pH tolerance based on the unique core-shell structure and the stability of polysaccharides against environmental stress. Furthermore, using the more acid-resistant phyA would maintain enzyme activity even after food processing and gastric digestion. Here, we report the preparation and optimization of core-shell hydrogels encapsulating phyA to provide thermal stability and environmental protection from salt and pH at high loading capacities while controlling particle size and water activity.

## 2. Materials and methods

### 2.1. Materials

Potassium chloride (KCl, > 99 %), alginic acid sodium salt from brown algae (medium viscosity), sodium hydroxide anhydrous (NaOH, > 97 %), calcium chloride dihydrate (CaCl_2_, > 99.5 %), ammonium molybdate tetrahydrate, trichloroacetic acid (TCA), fluorescein isothiocyanate isomer I (suitable for protein labeling, > 90 %), and L-ascorbic acid were purchased from Sigma-Aldrich (St. Louis, MO, US). Phytase from *Aspergillus niger* was supplied by DSM company (Heerlen, Netherlands). Pierce^TM^ rapid gold BCA (bicinchoninic acid) protein assay kits were purchased from Thermo Fisher Scientific (Liverpool, NY, US). Sodium phytate (95 %) was purchased from Astatech Inc. (Bristol, PA, US). κ-Carrageenan was purchased from TIC GUMS (Westchester, IL, US). Chitosan from *Aspergillus niger* was purchased from Sarchem Laboratories (Farmingdale, NJ, US). Sulfuric acid (95-98 %) was purchased from Fluka Honeywell (Charlotte, NC, US). Pepsin from pig gastric mucosa (∼2500 units/mg) was purchased from Roche (Basel, Switzerland).

### 2.2. Preparation of core-shell hydrogel beads

A 10.0 w/v% chitosan stock solution was made in distilled water. Its pH was adjusted to 5.0 using 0.2 M NaOH solution. The phyA stock solution (30.0 w/v%) and chitosan solution were mixed and diluted to result in three phyA concentrations (2, 4, and 16 w/v%) with one chitosan concentration (4.0 w/v%) using distilled water. These mixtures are vortexed for 1 minute at room temperature (∼25 ℃). The cross-linking agents, 0.2 M CaCl_2_ and 0.05 M KCl were dissolved in the phyA-chitosan solution to induce the complexation of phyA and chitosan, stirring for 30 min at room temperature.

Meanwhile, the alginate-κ-carrageenan solution was prepared by dissolving 0.5 g of sodium alginate and 0.5 g of κ-carrageenan in 100 mL of distilled water while stirring overnight at 60 ℃. After cooling to room temperature, the pH of the alginate-κ-carrageenan solution was adjusted to 10.0 using a 1.0 M NaOH solution.

To create the core-shell beads, the phyA-chitosan complex solution was loaded into a syringe and added slowly into the alginate-κ-carrageenan solution. Three different-sized needles were used to control the size of the droplets (Diameter: 18G-1.2 mm, 20G-0.9 mm, 25G-0.5 mm). Following the addition, the core-shell hydrogel beads are isolated from alginate-κ-carrageenan solution by centrifuge (6,700*g*) and stored at 4 ℃.

### 2.3. Encapsulation efficiency, released phytase, and loading capacity

We used a Pierce rapid gold BCA protein assay kit to determine how much phyA was incorporated into the core-shell hydrogel bead (Steć et al., 2023). The core-shell hydrogel beads were isolated by centrifugation at 6,700*g* for 10 min. The resulting supernatant (20 μL), containing the unencapsulated phyA, was treated with a 200 μL mixture of cupric sulfate and a copper chelator in a microplate well and incubated for 5 min at room temperature. The absorbance of the treated supernatant was measured at 480 nm using a plate reader (SpectraMax iD3, Molecular Devices, CA, US) to determine the quantity of phyA that was not encapsulated, free phytase; from that data, encapsulation efficiency, and loading capacity can be calculated. The encapsulation efficiency was calculated as the following equation (**Eq.1**):

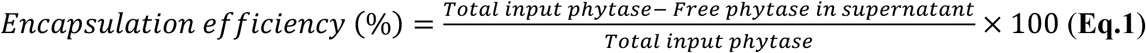

The loading capacity was calculated from the dehydrated mass and the loaded amount of phyA as the following equation (**Eq.2**):

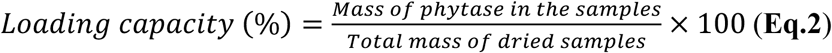

### 2.4. Phytase activity assay

A modified version of a previously reported method (Duru Kamaci & Peksel, 2021) was used to assess the encapsulated phyA activity. Briefly, 150 μL of substrate solution containing 0.44 mM sodium phytate in 100 mM sodium acetate buffer (pH 5.0) was pre-heated for 10 min at 50 °C in a water bath. One bead (approximately 150 μL) containing phyA was added to the substrate. The enzymatic reaction was allowed to occur for 30 min at 50 °C, after which time it was quenched by adding a denaturing agent (400 μL of 15 w/v% trichloroacetic acid (TCA) solution). After cooling the solution was centrifuged (6,700*g* for 10 min), then 100 μL of the resulting supernatant was added to a solution of 1 mL of the color reagent, a mixture of 10 w/v% ascorbic acid, 2.5 w/v% ammonium molybdate, and 1 M sulfuric acid solution at a ratio of 1:1:3 (v/v/v), and the entire solution was diluted to 900 μL using distilled water. This mixture was incubated for 15 min at 50 °C before cooling to room temperature for 10 min. The absorbance of the mixture was measured at 820 nm using a UV–Vis spectrophotometer (UV-2600, Shimadzu, Kyoto, JP). We used a standard curve based on phyA solution to determine the residual activity and thermal stability. In the case of thermal stability, the residual activity was measured after heating for 12 min at 100 °C.

### 2.5. Scanning electron microscope (SEM)

The microstructure of the samples was observed using a Zeiss Gemini 500 scanning electron microscope (Zeiss, DE). The samples were positioned on a carbon-taped stub and coated with gold using a sputter coater (Denton Desk V, NJ, US). Subsequently, the resulting samples were scanned and imaged using a secondary electron detector with a 20.0 μm aperture (accelerating voltage = 1 kV). After capturing, we calculated the average pore size using image J.

### 2.6. Dehydration process

After the sample preparation, the core-shell beads were dehydrated by air or vacuum. For air drying, wet beads were allowed to dry at 4 °C for 24 h or 50 °C for 2 h. Vacuum drying was done by heating at 30 °C for 1 h under vacuum conditions using a vacuum oven (VO 200, Memmert, Queens, NY, US). We measured the mass of samples before and after dehydration to calculate the water content, that is, the amount of bound water and free water in a sample. The water content was calculated as the following equation (**Eq.3**):

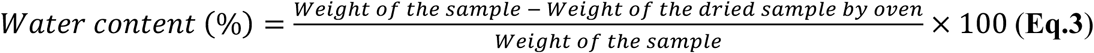

We also measured the water activity, that is, the energy status of free water as a parameter of the dehydrated sample, using a water activity meter (AQUALAB 4TE, Addium Inc., Pullman, WA, US). The water content was calculated as the following equation (**Eq.4**):

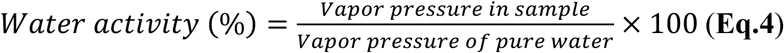

### 2.7. In vitro digestion

In vitro digestion was investigated according to a previously described method with slight modifications (Brodkorb et al., 2019). Simulated gastric fluid (SGF) was made by dissolving the proteolytic enzyme, pepsin 2000 U/mL, in a pH 2 buffer, adjusted by 2M HCl. The sample and free phyA with SGF were incubated at 37 ℃ for 2 h. After diluting with a pH 5.0 buffer, the residual activity of phyA was measured at pH 5.0 using the phytase activity assays.

### 2.8. Statistical analysis

Data are presented as means and standard deviations of triplicates and analyzed using Analysis of Variance (ANOVA). The Tukey HSD comparison test evaluated the differences between mean values (p < 0.05). All statistical analyses were performed using JMP Pro17 (SAS Institute, US) and plotted by GraphPad Prism10 (GraphPad Software Inc., US).

## 3. Results and Discussion

### 3.1. Optimization of phyA-loaded core-shell hydrogel beads

#### 3.1.1. Preparation of phyA-loaded core-shell hydrogel beads

We used a core-shell hydrogel bead system to encapsulate phytase from *Aspergillus niger* (phyA) for high thermal stability and acid resistance. We confirmed the phyA’s acid resistance by measuring the phyA activity under three pH conditions: pH 5 after incubation at pH 2 for 2 h, and at pH 5 after incubation at pH 2 for 2 h. **Figure 1(a)** We found that phyA activity can be recovered at pH 5 after being damaged by acidic conditions and that phyA has acid resistance (**Figure 1(a)**).

**Figure 1.**
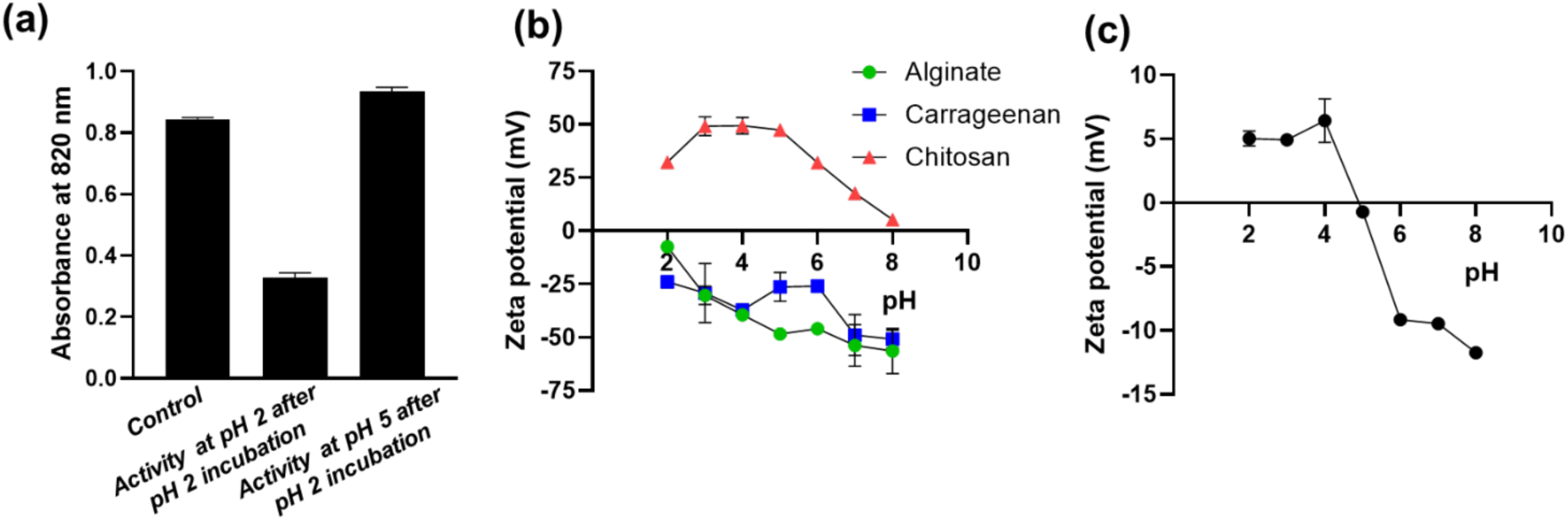
(a) Enzyme activity of phyA depending on the pH conditions. Zeta potential of (b) polysaccharides (alginate, carrageenan, and chitosan) and (c) phyA according to pH conditions.

To prepare the core complex between phyA and chitosan, because chitosan is positively charged below pH 6.5, the pH for complexation must allow phyA to be negatively charged. In the case of the polysaccharides used, chitosan is positively charged, while alginate and carrageenan are negatively charged in the applied pH conditions (**Figure 1(b)**). The zeta potential of phyA was measured to confirm the surface charge of phyA at different pH conditions (**Figure 1(c)**). The isoelectric point of phyA is between pH 4 and 5, so it has a slightly negative charge at pH 5. Based on this result, we chose pH 5.0 as the initial pH condition to induce the complexation of phyA and chitosan through electrostatic interactions.

The initial composition of core-shell hydrogel beads was a core of chitosan (4.0 w/v%) and phyA (2.0 w/v%) and a wall made from cross-linking alginate-κ-carrageenan (1:1, 1.0 w/v%) using calcium chloride (0.2 M) and potassium chloride (0.05 M) (E. Yang et al., 2024). The phyA-chitosan complex was formed when phyA was added to the chitosan solution at pH 5, indicated by the clear solutions becoming turbid. Then, the cross-linkers, CaCl_2_ and KCl, were added to the turbid solution, which was then added dropwise to the alginate-κ-carrageenan solution to form beads (**Figure 2(a)**).

**Figure 2.**
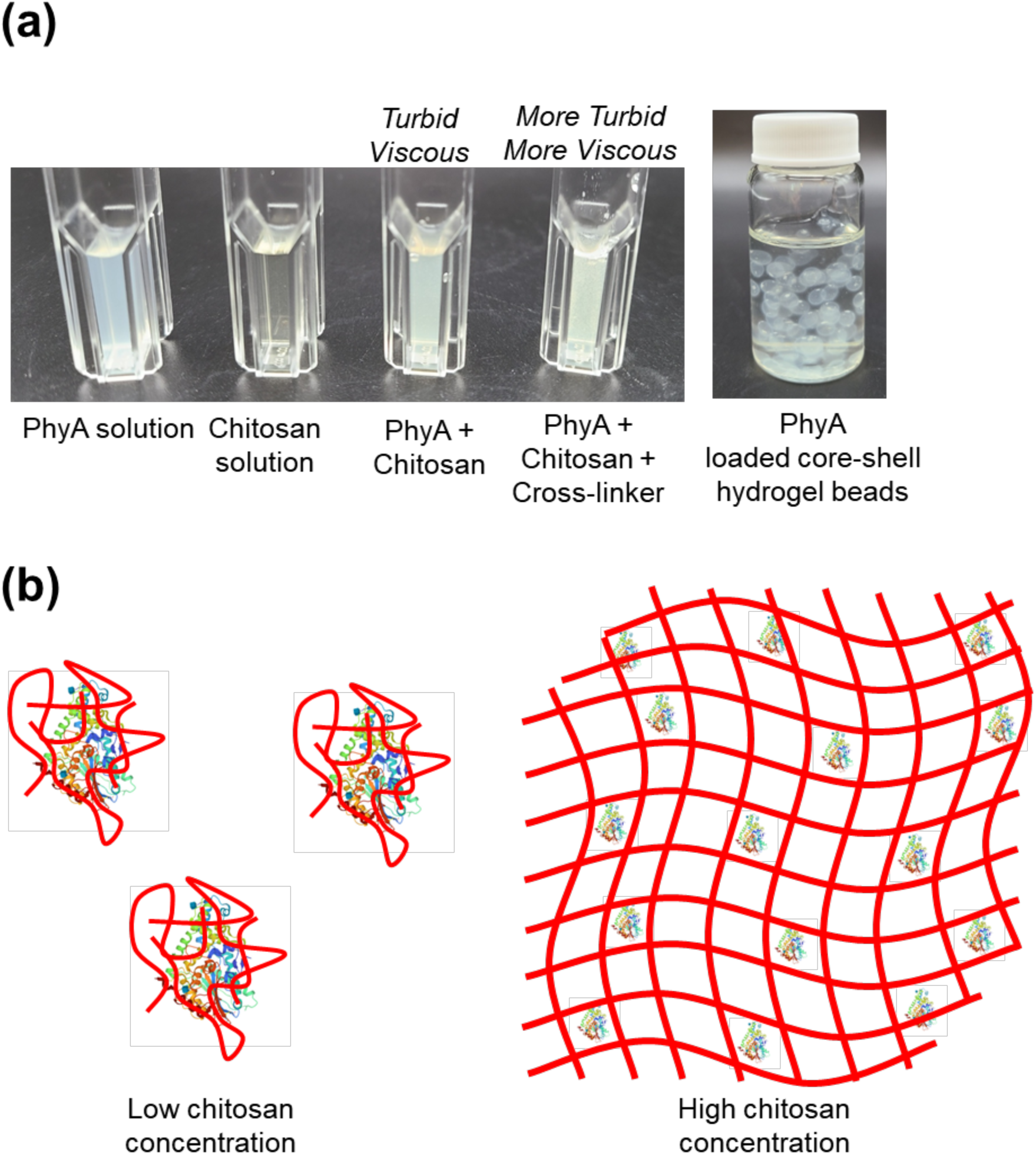
(a) Optical images of phyA, chitosan, phyA-chitosan complex with and without cross-linkers, and phyA-loaded core-shell hydrogel beads. (b) Schematic illustration of phyA-chitosan complex depending on the chitosan concentration.

The initial turbidity and viscosity of the phyA-chitosan solution indicate that a protein-polysaccharide hydrogel has formed (Ma et al., 2018). Depending on the ratio of phyA and chitosan, the hydrogel structure (**Figure 2(b)**) was calculated using the theoretical size of chitosan (molecular weight: 200kDa, expected length (fully solubilized): ∼300 nm (Hailei et al., 2016)) and phyA (molecular weight: 99kDa, expected size: ∼6 nm (Oakley, 2010)). After mixing with the CaCl_2_ and KCl, the solution became more turbid and viscous. It is likely that the increased ionic strength of the solution caused the structure of the phyA-chitosan complex to contract due to the formation of additional network interactions (J. Yang et al., 2019). Usually, chitosan gelation is induced by an anionic cross-liker like TPP (tripolyphosphate) or genipin. In the case of a protein-polysaccharide hydrogel, however, the addition of calcium ions enhances the hydrophobic interaction and increases the hydrogen bonds between the enzyme and the polysaccharide, but may have decreased electrostatic interactions between them because of the increased ionic strength (Yan et al., 2022). Using this highly viscous solution, we prepared uniformly-sized white phyA-loaded core-shell beads with high encapsulation efficiency (94.0 %) and low loading capacity (20.6 %). Then, based on this result, we optimized the preparation conditions to enhance loading capacity and control bead size, retaining thermal stability for practical application in the food industry and animal feeding.

#### 3.1.2. Loading capacity of core-shell hydrogel beads

For practical application in the food industry, our goal was to encapsulate as much phyA as possible with the highest loading capacity while balancing the thermal stability of the protein and recognizing that increasing the phyA concentration with respect to chitosan concentration would mean that it was less well protected (**Table 1**). We calculated the maximum phyA concentration that chitosan can cover to maintain thermal stability and increased the phyA ratio in the phyA-chitosan complex core to improve loading capacity. Two kinds of core-shell beads, chitosan 4 w/v%-phyA 2 w/v% and chitosan 4 w/v%-phyA 16 w/v%, were prepared. As a result, the phyA 16 w/v% sample has a 2.5 times higher loading capacity (52.2 ± 2.4 %) than the phyA 2 w/v% sample (20.6 ± 1.3 %).

**Table 1.** Loading capacity, encapsulation efficiency, and thermal stability of phytase A-loaded core-shell hydrogel beads depending on the phytase concentration.

| **PhyA concentration** | **Loading capacity (%)** | **Encapsulation efficiency (%)** | **Thermal stability (%)** |
| --- | --- | --- | --- |
| 2 w/v% | 20.6 ± 1.3 | 94.0 ± 3.4 | 83.9 ± 3.5 |
| 16 w/v% | 52.2 ± 2.4 | 82.6 ± 3.4 | 69.4 ± 9.5 |

The encapsulation efficiency of the phyA 16 w/v% (82.6 ± 3.4 %) sample was 11.4% lower than that of the phyA 2 w/v% sample (94.0 ± 3.4 %). The thermal stability of phyA in these two core-shell beads was also measured, which confirmed that at higher phyA concentration (16 w/v%), there was reduced thermal stability (69.4 ± 9.5 %) compared to phyA 2 w/v% (83.9 ± 3.5 %). The decrease in thermal stability was much smaller than the increase in loading capacity, and the thermal stability was similar once the phyA concentration exceeded 8 w/v% (**Figure S1**).

These results can be attributed to three possible interactions. First, phyA in the complex can act like the protein composite in the protein-polysaccharide hydrogel, such as WPI-chitosan hydrogel (Liu et al., 2020) and zein-chitosan hydrogel (Ma et al., 2018). Based on the positively charged amino groups, various interactions between protein and chitosan are possible, including electrostatic interactions and hydrogen bonding. Thus, phyA can enhance the core structure by forming densely branched networks of chitosan chains. A protein composite would then require less chitosan to cover the rest of the protein because it would be partially covered by the chitosan in the composite.

Second, the dense network created by the composite would increase the viscosity of the initial droplet. The high viscoelasticity of the droplets helps maintain the droplet shape while it falls into an alginate-κ-carrageenan solution and eventually increases encapsulation efficiency. Finally, compared with other hydrogels, the core-shell hydrogel has a shell structure that maintains encapsulation efficiency and protects the core, regardless of the core composition. For example (Zhang et al., 2016), alginate hydrogel beads have a low encapsulation efficiency (∼20%) for protein at the same pH condition, pH 5, because of the electrostatic repulsion between protein-alginate. Therefore, increasing phyA concentration to 16 w/v% significantly improves the loading capacity while maintaining acceptable thermal stability of the phyA-loaded beads.

#### 3.1.3. Size control of core-shell hydrogel beads

Particle/bead size control is important for mouthfeel when incorporated into foods and beverages and to eliminate the need to chew the beads. Shear stress or compression caused by chewing can cause fragmentation, which is the physical disruption of the carriers that induces active materials to be released quickly and would lead to loss of thermal stability of the protein and protection from very low pH. The average size of the food sample is 2 mm after chopping by chewing; therefore, it is at the mouth. This means that the particle or bead size should be smaller than 2 mm (Kim et al., 2022).

There are several factors that can be adjusted to control the size of hydrogel beads, including preparation method and cross-linker concentration. We used reverse gelation to form our core-shell hydrogel beads, which is the fabrication of liquid core beads through reverse extrusion dropping. Compared to conventional extrusion by syringe, which produces large gel particle sizes, this method makes smaller-sized beads (Saqib et al., 2022). Because the size of alginate hydrogel beads is controlled by CaCl_2_ concentration, we used a higher CaCl_2_ concentration to make a smaller-sized bead. In the case of core-shell beads, the CaCl_2_ concentration controls the alginate shell thickness. Unfortunately, we found that recovery of phyA activity and thermal stability differed depending on the shell thickness, so to control the core size, the initial size of the droplet became an essential factor.

To make smaller-sized droplets, we adjusted the needle diameter, the smallness of which was limited by the viscosity (> 200 cP), which caused pumping difficulties and needle blockage when the needle was too small. We prepared the samples using three-needle diameters to control the bead size, and as expected, the beads became smaller using a smaller needle (**Figure 3(a)**). The smallest bead size using a syringe needle without clogging was 2.5 mm, slightly bigger than our target size of 2 mm, though after dehydration, we should see additional shrinkage and get closer to our target (Wong et al., 2021).

**Figure 3.**
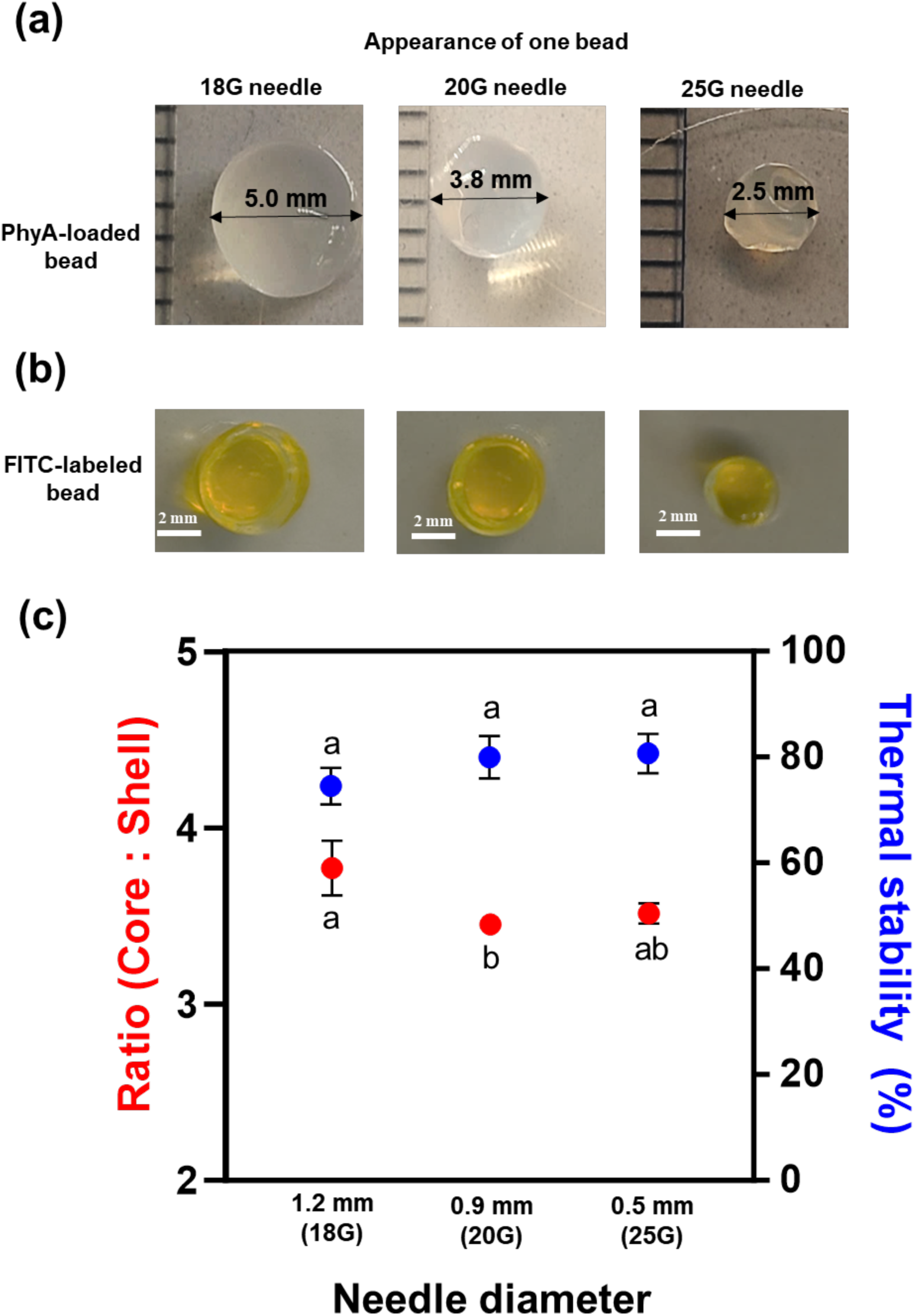
Optical image of (a) phyA-loaded and (b) FITC-labeled core-shell hydrogel beads depending on the needle diameter. (c) The thickness ratio of core and shell and thermal stability of phyA-loaded core-shell hydrogel beads according to needle diameter. A connecting letter plot was used at the significance difference level of p ≤ 0.05.

The thickness ratio between the core and the shell is more important for the protective effect than the thickness of the shell. Fluorescein isothiocyanate (FITC) is used to visualize the core and the shell and calculate the thickness ratio of the core and shell. The thickness of the shell is determined by the initial concentration and amount of the diffused crosslinker. Once the shell is formed, the diffusion of the crosslinker becomes restricted, making it difficult for the thickness to increase further with reaction time. Additionally, the amount of crosslinker initially diffusing is determined by the surface area of the droplet in contact with the alginate-carrageenan solution. Therefore, the diameter of the needle used to control the droplet size has a significant impact. As a result, the core and shell thickness are dependent on the needle diameter (**Figure 3(b)**). Also, the thickness ratio and thermal stability were inversely proportional (**Figure 3(c)**)

The larger the ratio of the shell to the core containing the phytase to be protected, the more the phytase is physically shielded from heat and benefits from the protective function of the shell. Consequently, the optimum condition for making the smallest bead with high thermal stability is 0.5 mm needle diameter.

### 3.2. Effect of the dehydration process

In the food industry, controlling water activity is a critical issue for the broad application and storage stability to inhibit microorganism growth. In the case of hydrogel beads, because of their unique structure that holds a large amount of water, dehydration can do one of two things: (1) If the gel structure of the shell is maintained during dehydration, the gel structure becomes tighter as water molecules disappear, which will better protect the phyA from heat. (2) If the shell structure is broken during dehydration, the phyA will be released, and the hydrogel beads will lose the protective effect (Kondaveeti et al., 2022; Wong et al., 2021).

Compared to normal hydrogel beads, the dehydration process of core-shell hydrogel beads has two phases with different dehydration rates because of the core-shell structure. To find a proper dehydration method, various drying methods were investigated, such as freeze-drying, spray-drying, air-drying, and vacuum-drying. During freeze-drying (E. Yang et al., 2021), ice crystallization and ice crystals formed that damaged the shell network. In the case of spray-drying, the gel network was made permeable during the heating process at 170-180°C (J. Wang et al., 2023). Therefore, vacuum drying and air drying at mild temperatures, lower than the degradation temperature of the components of the shell, gave acceptable results (Sánchez-Fernández et al., 2021; Zhang et al., 2016).

Our goal was to reduce the water activity below a_w_ 0.6 to control the microorganism growth. Three conditions were investigated: vacuum drying at 30 °C for 1 h, air drying at 50 °C for 2 h, and air drying at 4 °C for 12 h. In the case of vacuum drying, we compared incubation temperatures (30, 60, and 90 °C) to select mild conditions (**Figure 4**). As a result, we selected 30 °C because it took 60 min to get 2 w/w% water content with a_w_ 0.35 water activity. Air-dried samples (50 °C for 2 h and 4 °C for 12 h) also have around 2 w/w% water content after dehydration.

**Figure 4.**
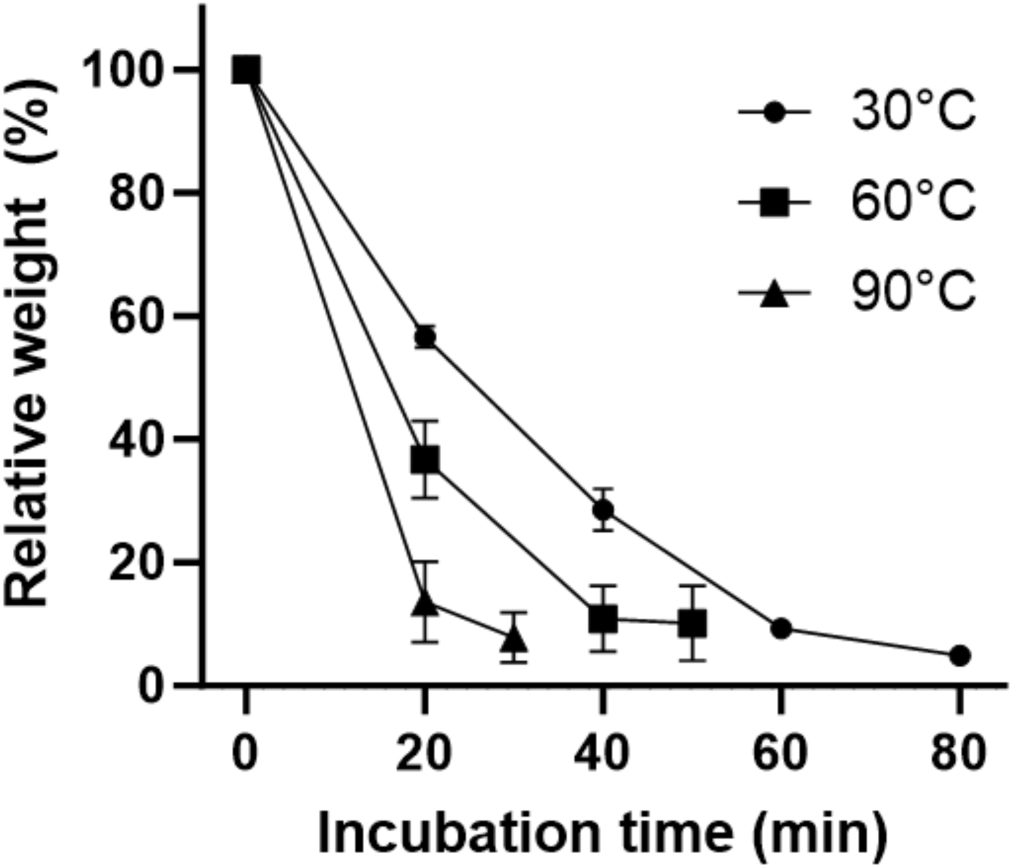
Effect of temperature on the relative weight of hydrogel beads during vacuum drying.

The optical images of the dehydrated samples using the different dehydration conditions and rehydrated samples for 15 and 30 min were used to gauge the impact of dehydration (**Figure 5(a)**). The bead that was air-dried at 4 °C was rehydrated for 15 min and showed the same morphology as before drying. The bead incubated at 50 °C did not fully recover its former morphology even after rehydration for 30 min.

**Figure 5.**
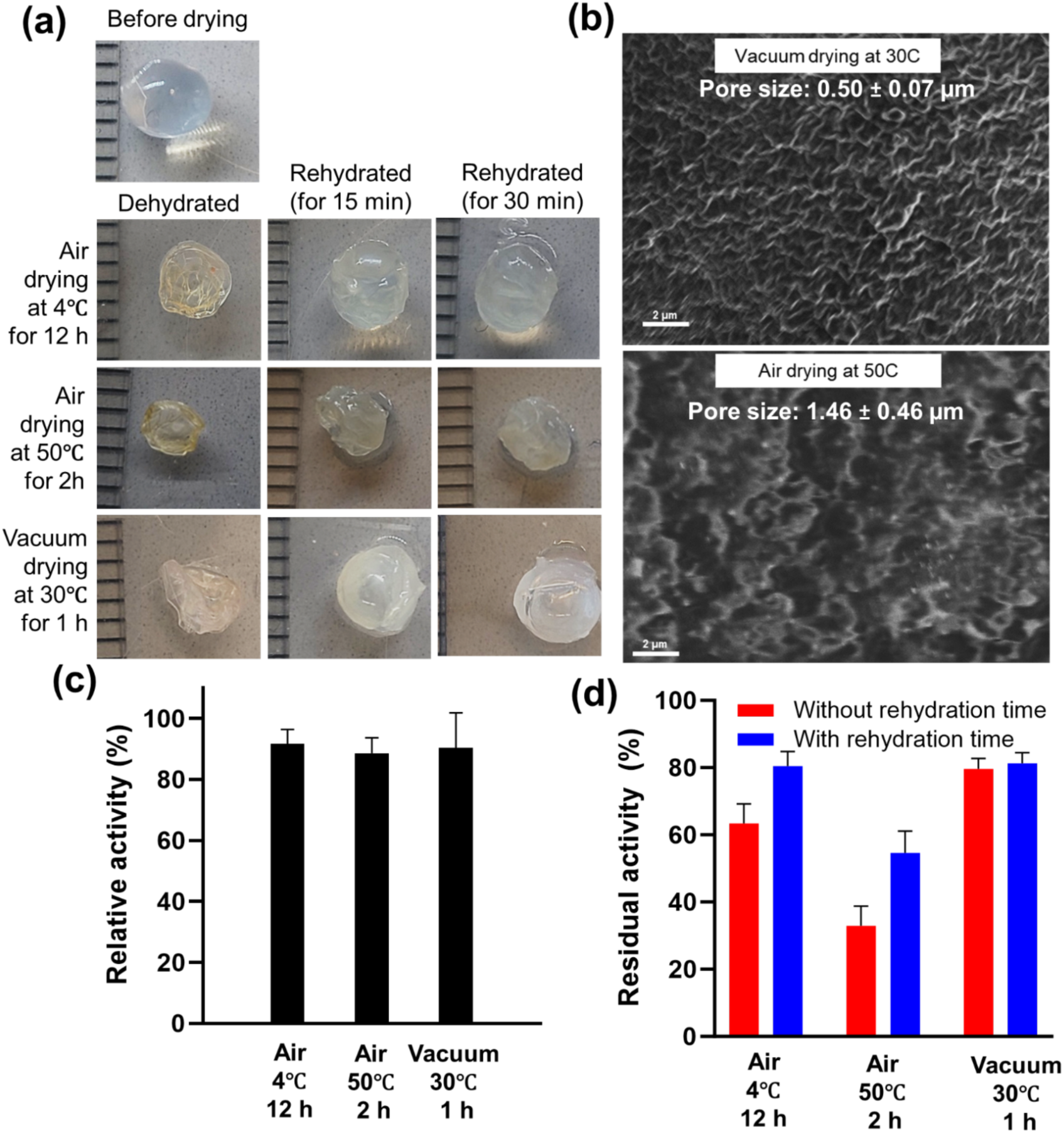
(a) Optical images of core-shell hydrogel beads after dehydration and rehydration. (b) SEM images of core-shell hydrogel beads after dehydration. (c) Relative activity of core-shell hydrogel beads after dehydration and rehydration. (d) Thermal stability of core-shell hydrogel beads after dehydration with or without rehydration process.

Further, a comparison of the surface morphology by SEM showed that the gel network of the air-dried bead at 50 °C was partially broken (**Figure 5(b)**). When we calculated the pore size using image J, the pore size of the 50 °C air-dried sample was 1.46 ± 0.46 µm, which was much bigger than the vacuum-dried sample (0.50 ± 0.07 µm). Unlike air-drying, the gel network of the vacuum-dried sample was maintained after dehydration, indicating that the shell integrity is important to the recovery of beads after dehydration and rehydration (Kondaveeti et al., 2022).

The phyA activity and thermal stability after dehydration were measured (**Figure 5(c)**). Each sample retained enzyme activity between 88.6 to 91.7%, indicating that air or vacuum drying does not affect the phyA-chitosan composite.

In the case of thermal stability, however, which is the residual activity after heating at 100 °C for 12 min (**Figure 5(d)**), there are significant differences, depending on the dehydration condition. When the activity was measured for the dehydrated samples, vacuum drying protected almost 80% of activity (79.7 ± 3.0 %), but air drying at 50 °C lowered the activity more substantively (33.0 ± 5.9 %). This is consistent with the loosening of the gel network of the shell, resulting in the exposure of the phyA to heat stress. When the beads were rehydrated before heat treatment at 100 °C for 12 min, the phyA activity was higher after heat treatment than for the dehydrated beads. For example, the air-dried sample at 4 °C has 17% higher thermal stability after rehydration (80.5 ± 4.3 %) than the dry bead (63.4 ± 5.8 %). The rehydration process makes the dehydrated samples swell. As a result, the beads with water take more time to reach equilibrium temperatures than those without water, resulting in enhanced thermal stability. The optimum condition to retain the most phyA activity is to vacuum-dry at 30 °C for 1 h and then, before use, rehydrate for 30 min before heating the beads.

### 3.3. Effect of environmental stress

When the environmental conditions are changed, there are several possibilities for releasing the cargo from hydrogel beads, such as diffusion, swelling, shrinking, fragmentation, and erosion (Saqib et al., 2022). Diffusion is the most commonly occurring material release process. The diffusion rate depends on the osmotic pressure, pore size, size of the core material, pH, and temperature. In the case of core-shell hydrogel, diffusion is controlled by the shell’s properties, such as its thickness and porous structure. In this process, the encapsulated phytase can be diffused from the core to the surface of the shell matrix and then dispersed and transported by the surrounding medium.

#### 3.3.1. Effect of temperature

Enzyme encapsulation can mitigate the effect of temperature so that encapsulated enzymes have higher thermal stability than soluble unencapsulated enzymes (Karim et al., 2017). Using the optimized sample (16 w/v% phyA:4 w/v% Chitosan, vacuum-dried at 50 ℃ and rehydrated for 30 min), we investigated the effect of temperature on the activity of the encapsulated phyA. The preparation of plant-based foods, such as maize porridge and bouillon, requires boiling in water. For example, maize porridge is heated for more than 5 min at 100 ℃, and bouillon is made by heating for 30 min at 90 ℃ (S. Wang et al., 2024). To assess the impact of food preparation conditions on our core-shell hydrogel beads, we incubated our beads at room temperature, 50 ℃, and 100 ℃ for 60 min.

Diffusion of core material and surface erosion are expected as the release mechanism caused by high temperatures. First, increasing the temperature makes the diffusion rate of the core material high, causing it to expand while at the same time, the shell weakens and becomes more porous (van den Berg et al., 2022). This kind of release pattern usually follows a linear relationship. During shell erosion due to the breakdown of the gel network, heat can be withdrawn from the surface to the core gradually or entirely. After breaking bonds, the external fluid enters the food matrix, transporting the encapsulated phytase into the surrounding liquid system. The erosion rate depends on the physicochemical stability of the food matrix, the molecular weight or gelling mechanism of biopolymers, or the efficacy of the release medium.

The released amount of phyA (**Figure 6(a)**) and relative activity were measured during incubation (**Figure 6(b)**). The release at room temperature is lower than 10% after 1 h, based on the simple diffusion. Incubation at 100 ℃, however, shows that phytA is released in direct proportion to heating time, while the release rate at 50 ℃ is slow for the first 40 min (17.6 ± 4.2 %) and increases over the next 40-60 min (35.5 ± 10.0 %). At 50 ℃, the release is initially controlled by diffusion, and then, as time passes, the release becomes a combination of diffusion and erosion.

**Figure 6.**
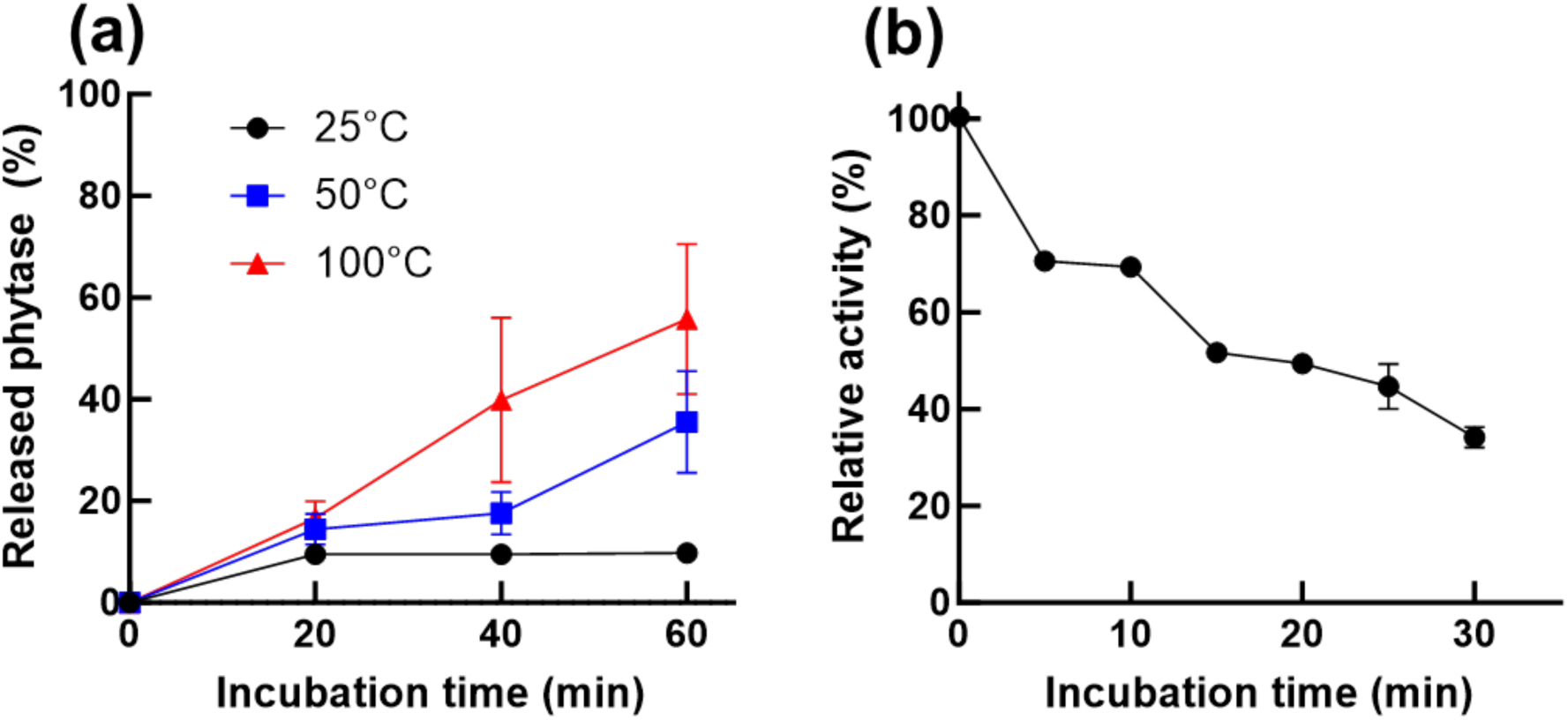
Effect of temperature on (a) release profile at 25 °C, 50 °C, and 100 °C and (b) residual activity of phytase A-loaded core-shell hydrogel beads at 100 °C.

This core-shell hydrogel bead formulation has a 90 °C degradation temperature based on the complex degradation of the core and shell matrix (E. Yang et al., 2024), Another study (Fajardo et al., 2012) reported that the degradation temperature (258 °C) of the alginate–Ca^2+^ matrix is associated with the destruction of glycosidic bonds between the glucose backbone of alginate. Alternatively, a κ-carrageenan xerogel has a much lower degradation temperature (71.2 °C) because of its heat-inducible transformation (Zhang et al., 2016). Because of these examples, we expected that heating higher than 90 °C or long-term heating would first break down the shell’s κ-carrageenan gel network, resulting in partial erosion followed by diffusion.

After 15 minutes, the encapsulated phyA retained 51.7 ± 1.5 % of its activity at 100 °C (**Figure 6(b)**). The activity half-life is around 17 min at 100 °C, which corresponds to higher thermal stability than raw phyA (27.0% of activity at 100 ℃ for 12 min). This means that core-shell hydrogel beads protect phyA’s activity against extreme heating conditions. Based on the release profile, the activity loss is higher than the released amount of phyA after 60 minutes because the still encapsulated phyA is also exposed to heat due to the partially broken shell.

#### 3.3.2. Effect of salt concentration on the phytase activity

The salt type and concentration were decided based on the conditions in the food matrix and digestion model. As a model food matrix, the composition of maize porridge contains sodium chloride with a concentration of around 0.24 w/v% (41 mM). Also, calcium and bile salts are present in the human gastrointestinal tract. Following the INFOGEST digestion model (Brodkorb et al., 2019), there are several kinds of salts, including calcium 0.5 w/v% (∼1.5 mM) and bile salts (10 mM) in simulated digestion fluid.

The amount of phyA released after incubation with sodium chloride 50-300 mM solution for 2 h increased with increasing concentration (**Figure 7(a)**). The concentration of cross-linkers in the initial chitosan-phyA solution is lower than their starting concentrations of 200 mM for the calcium ion and 50 mM for the potassium ion because some of them are bound to the alginate and k-carrageenan molecules during gelation. We attribute the high release of phyA at sodium ion concentrations of 300 mM to a shrinking phenomenon as the phyA is released with water due to osmotic pressure imbalance. As a result, there is about 43.8 ± 5.5 % release at 300 mM, much higher than at 200 mM (20.7 ± 2.3 %). In the case of phyA activity, even though the released phyA, there is only an 8% decrease of activity until 300 mM of sodium chloride (92.2 ± 4.7 %) (**Figure 7(b)**) because phyA has good salt tolerance itself (Sandberg et al., 1996). However, if heated after incubation with salt, activity loss will be similar to the released phyA.

**Figure 7.**
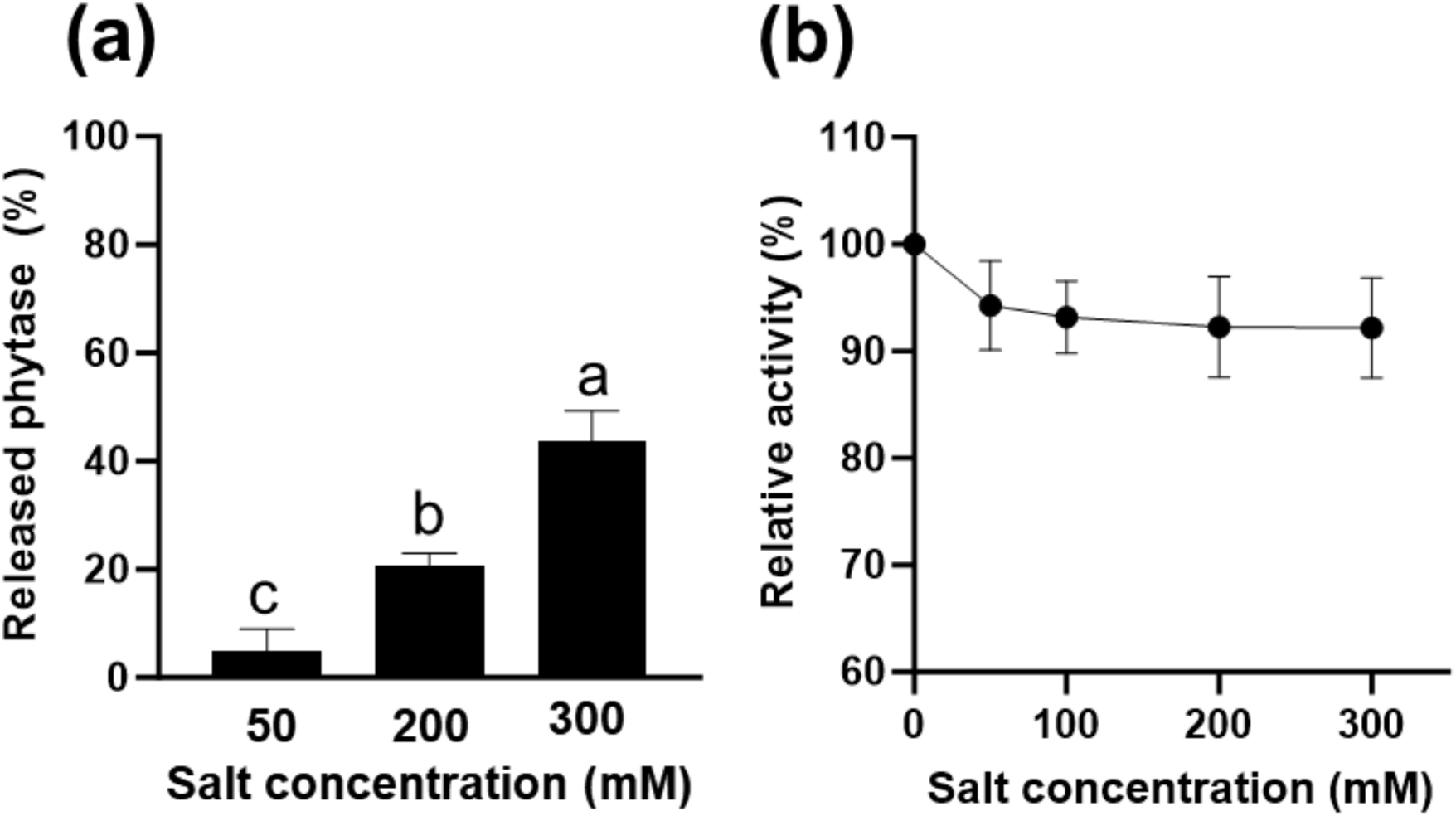
Effect of salt concentration of the phytase A (a) release and (b) relative activity in the core-shell hydrogel beads. A connecting letter plot was used at the significance difference level of p ≤ 0.05.

#### 3.3.3. Effect of pH conditions on the phytase activity

The effect of pH on enzymes and hydrogel beads is well-known (Lima et al., 2018). The targeted pH conditions were based on the food matrix and digestion conditions. Because the samples are intended for addition to maize porridge (pH 5-7) and the gastrointestinal tract, including a mouth (pH 7), gastric phase (pH 2), and small intestine (pH 7), we selected pH 2 and 7 as the representative pH conditions for release studies. We measured the released amount of phyA after incubation with pH 2 and 7 buffers for 30 and 120 min, respectively (**Figure 8(a)**). The final amount of protein released at the end of the incubation period (120 min) depended strongly on the pH: pH 2 (19.9 ± 0.9 %) and pH 7 (67.4 ± 0.7 %).

**Figure 8.**
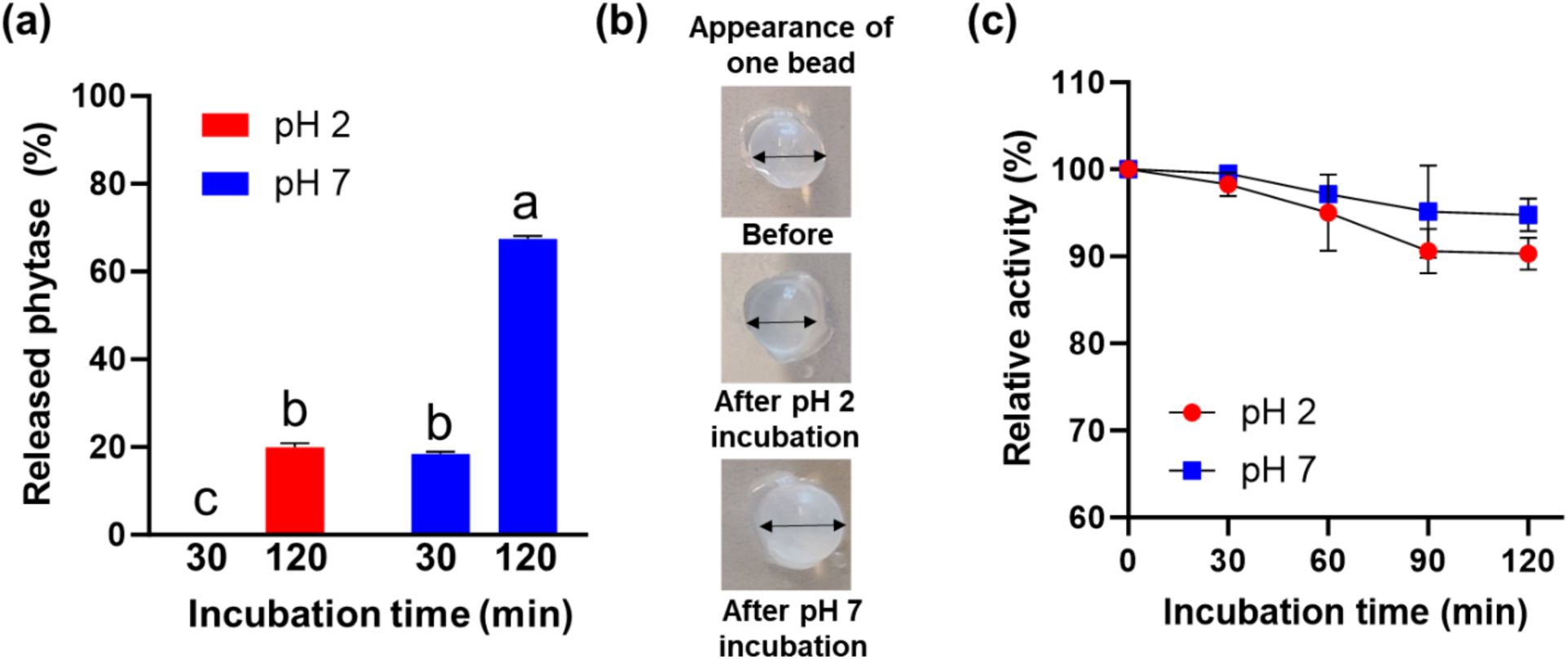
Effect of pH condition on the phytase A (a) release, (b) appearance, and (c) relative activity of phytase A-loaded core-shell hydrogel beads. A connecting letter plot was used at the significance difference level of p ≤ 0.05.

Variations in pore sizes in hydrogels respond differently to changes in pH (Lima et al., 2018; Zhang et al., 2016). Alginate normal hydrogels exhibit larger pores when swollen in basic pH compared to acidic pH. These differences in porosity relate to the presence of ionizable groups within the hydrogel network. At acidic pH, alginate’s carboxyl groups (with a pKa 4.5) exist in a non-ionized (-COOH) state, leading to stronger attractive forces that cause the polymer network to contract slightly, resulting in a denser structure that affects water absorption (**Figure 8(a)**-pH 2).

Combining alginate with κ-carrageenan forms a denser network that decreases the release of phyA at pH 2 compared to alginate-only hydrogels. Above pH 4.5, the carboxylic acid groups are deprotonated to form carboxylates, creating electrostatic repulsion between the anionic groups, which causes the polymer network to expand and the pores to increase, allowing the cargo to release (**Figure 8(a)**-pH 7). The pore size and gel network changes, influenced by environmental changes, contribute to the hydrogels’ morphological properties, as shown in their appearance (**Figure 8(b)**).

Compared to the initial beads that were formed, the bead pH 7 was swollen and opaque after incubation. This change can be attributed to differences in the electrostatic interactions of the chitosan-phyA and alginate-κ-carrageenan at different pH levels. From pH 2 to 5, phyA and chitosan are both positively charged, while alginate and κ-carrageenan have a negative charge. As the phyA diffuses, it will come into contact with the positively charged shell materials and will be strongly attracted to anionic polysaccharides on the surface of the bead. Conversely, at higher than pH 6, the phyA has a negative charge, and alginate and κ-carrageenan also have a negative charge. Like the simple alginate hydrogel, the protein and alginate would have similar charges and be electrostatically repulsed, and the protein would be excreted from the hydrogel network. Thus, the network had larger pores at pH 7, which partially resulted in an accelerated release of the phytase. These results indicate that phyA-loaded beads could release phyA, which is responsible for the observed results under basic pH conditions. This pH-responsive property makes the hydrogel system protect core materials in the pepsin of the stomach and then release them in the neutral environment of the small intestine.

Meanwhile, in the case of phytase activity (**Figure 8(c)**), even though the released phyA is 19.9 %, there is a 9.7 % decrease in activity (90.3 ± 1.8 %) in pH 2 after incubation for 2 h. This result is because phyA has good tolerance to acidic conditions, as mentioned before (Sandberg & Scheers, 2016). Therefore, encapsulated phytases can maintain activity continuously in both the stomach and the small intestine, making it much more advantageous for phytate hydrolysis before mineral absorption.

### 3.4. In vitro digestion

#### 3.4.1. Effect of pepsin on the activity of phytase

In food applications, the phytase-loaded beads must protect against thermal treatment during preparation as well as survive the digestive tract to the small intestine, where mineral absorption takes place in the duodenum. PhyA is also susceptible to proteolytic cleavage and inactivation by pepsin. To assess the core-shell beads’ ability to protect phyA from gastric juices, we measured the activity after treating gastric fluids, including pepsin and very low pH (**Figure 9(a)**). After heating at 100 °C for 12 min and the digestion simulation, the phyA in the beads has 62.0 ± 3.2 % activity. Compared with raw phyA, it is a promising result.

**Figure 9.**
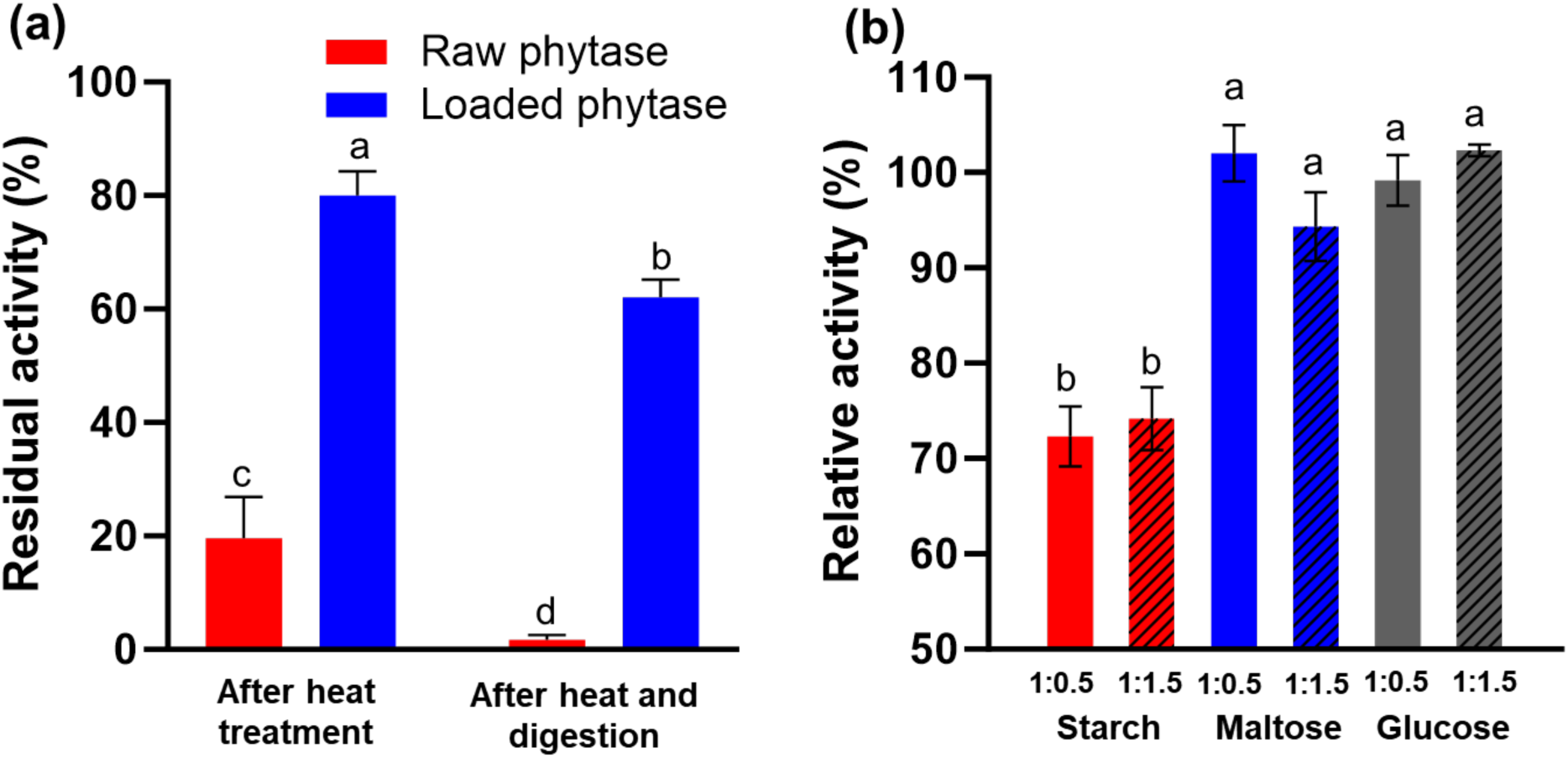
(a) Residual activity of phytase A-loaded core-shell hydrogel beads after heat and digestion. (b) Relative activity of phytase A in core-shell hydrogel beads with different substrate types, including starch, maltose, and glucose. Different letters indicate a significance difference level of p ≤ 0.05.

The protective effect against pepsin was expected based on this encapsulation system’s component. In the case of other encapsulation systems, including protein (Atma et al., 2024), pepsin accelerates the release of core materials. This system consists of three kinds of polysaccharides so that it can avoid the effect of pepsin. In particular, a recent study showed that the alginate-chitosan beads displayed excellent stability in gastric fluid (Wong et al., 2021). Furthermore, in our encapsulation system, the chitosan in the phyA-chitosan complex and the alginate-carrageenan shell both act as a barrier, blocking pepsin penetration and preventing phyA hydrolysis.

#### 3.4.2. Effect of food components

The major component of maize porridge is starch (about 75 w/w%), which can bind with other materials like phytate. As a preliminary test, we mixed phytate with starch and measured the average diameter using DLS analysis. Thephytate-starch complexes have 5 μm of average diameter (**Figure S2**). These are bigger than the pore size of the shell, 0.5 micrometers based on the SEM analysis, so they cannot penetrate the shells to impact the phyA in the core. If the starch is digested, however, into smaller polysaccharides, around 170 - 320 Da, we expect it to diffuse into the core.

We measured the activity of phyA in the beads after mixing phytate with saccharides such as starch, maltose, and glucose to investigate the effect of polysaccharides and glucose. The enzymatic activity using phytate with starch as substrate decreased by 26 - 28 % (1:0.5 (72.4 ± 3.2 %), 1:1.5 (74.2 ± 3.3 %)), while the activity in the presence of maltose was slightly reduced, and glucose had no impact on activity (**Figure 9(b)**). This means that starch in maize porridge affects phyA activity in the beads. Phytates with disaccharides and monosaccharides, however, can diffuse into the shell and be degraded.

## 4. Conclusions

A core-shell hydrogel bead was fabricated to maintain the catalytic activity of phyA and protect it from heat treatment and proteases. These hydrogel beads have a core composed of a phyA-chitosan complex that stabilizes and protects the phyA and a shell consisting of cross-linked alginate-κ-carrageenan that provides additional protection and heat resistance. We developed its property for food application and evaluated this system against environmental conditions. Optimization of the ratio of chitosan to phyA improved the loading capacity from 20.6% to 52.2% with high encapsulation efficiency (82.6%) and thermal stability of the phyA activity (69.4%). Reducing the needle diameter from 1.2 mm to 0.5 mm reduced the bead size to 2.5 mm. Dehydration of the beads by vacuum drying for 1 h at 30 °C protected the beads from shell damage with a uniform pore size (0.5 µm) so that the dehydrated sample can protect 79.7% enzyme activity against heating. As a result, the core-shell hydrogel beads showed good tolerance against temperature, salt, pH, and protease based on the structure combined with phyA-chitosan and alginate-carrageenan. Notably, compared with free phyA, loaded phyA can maintain 62% activity even after heat treatment at 100 °C for 12 min and gastric digestion. Regarding the food matrix, we found that phytate binding with glucose after digestion diffuses into the core-shell and is degraded by the phyA inside the beads. These results suggest that the core-shell beads are suitable for protecting phyA activity against various stressors, which may make them suitable for food applications.

## Generative Artificial Intelligence Statement

The authors certify that generative AI was not used in preparing this article. Non-generative AI, such as spelling and grammar checkers in Office 365 and Google Docs, and citation managing software, was used. All instances when non-generative AI was used were reviewed by the authors and editors.

## Data Availability Statement

All data used to produce the figures in this article and the SI are freely available at: https://doi.org/10.5281/zenodo.14920624.

## Author CRediT statement

**Eunhye Yang:** Conceptualization, investigation, formal analysis, visualization, writing-original draft, reviewing and editing. **Waritsara Khongkomolsakul:** Conceptualization, investigation, writing review, and editing. **Younas Dadmohammadi:** Project conceptualization, supervision, funding acquisition, writing review, and editing. **Alireza Abbaspourrad:** Project conceptualization and administration, supervision, funding acquisition, writing review, and editing

## Acknowledgments

This work was carried out with the support of the Nutrition R&D Program (INV-043930) provided by the Bill & Melinda Gates Foundation (WA, US). The SEM facility from the Cornell Center for Materials Research (CCMR) was supported by the National Science Foundation under Award (DMR-1719875).

## References

1. Brodkorb, A., Egger, L., Alminger, M., Alvito, P., Assunção, R., Ballance, S., Bohn, T., Bourlieu-Lacanal, C., Boutrou, R., Carrière, F., Clemente, A., Corredig, M., Dupont, D., Dufour, C., Edwards, C., Golding, M., Karakaya, S., Kirkhus, B., Le Feunteun, S., … Recio, I. (2019). INFOGEST static in vitro simulation of gastrointestinal food digestion. Nature Protocols, 14(4), 991–1014. 10.1038/s41596-018-0119-1

2. Brouns, F. (2022). Phytic Acid and Whole Grains for Health Controversy. Nutrients, 14(1). 10.3390/nu14010025

3. Duru Kamaci, U., & Peksel, A. (2021). Enhanced Catalytic Activity of Immobilized Phytase into Polyvinyl Alcohol-Sodium Alginate Based Electrospun Nanofibers. Catalysis Letters, 151(3), 821–831. 10.1007/s10562-020-03339-0

4. Fajardo, A. R., Silva, M. B., Lopes, L. C., Piai, J. F., Rubira, A. F., & Muniz, E. C. (2012). Hydrogel based on an alginate–Ca2+/chondroitin sulfate matrix as a potential colon-specific drug delivery system. RSC Adv., 2(29), 11095–11103. 10.1039/C2RA20785K

5. Hailei, Z., Yinli, L., Xu, Z., Bo, L., Huiling, Z., & Dong, C. (2016). Directly determining the molecular weight of chitosan with atomic force microscopy. Front Nanosci Nanotech. 10.15761/FNN.1000121

6. Karim, A., Nawaz, M. A., Aman, A., & Ul Qader, S. A. (2017). Role of Anionic Polysaccharide (Alginate) on Activity, Stability and Recycling Efficiency of Bacterial Endo (1→4) β-d-Glucanase of GH12 Family. Catalysis Letters, 147(7), 1792–1801. 10.1007/s10562-017-2074-9

7. Kim, E. H.-J., Wilson, A. J., Motoi, L., Mishra, S., Monro, J., Parkar, S. G., Rosendale, D., Stoklosinski, H. M., Jobsis, C. M. H., Wadamori, Y., Hedderley, D. I., & Morgenstern, M. P. (2022). Chewing differences in consumers affect the digestion and colonic fermentation outcomes: In vitro studies. Food & Function, 13(18), 9355–9371. 10.1039/D1FO04364A

8. Kondaveeti, S., Petri, D. F. S., & Jeong, H. E. (2022). Efficiency of air-dried and freeze-dried alginate/xanthan beads in batch, recirculating and column adsorption processes. International Journal of Biological Macromolecules, 204, 345–355. 10.1016/j.ijbiomac.2022.02.011

9. Lima, D. S., Tenório-Neto, E. T., Lima-Tenório, M. K., Guilherme, M. R., Scariot, D. B., Nakamura, C. V., Muniz, E. C., & Rubira, A. F. (2018). pH-responsive alginate-based hydrogels for protein delivery. Journal of Molecular Liquids, 262, 29–36. 10.1016/j.molliq.2018.04.002

10. Liu, Z., Liu, C., Sun, X., Zhang, S., Yuan, Y., Wang, D., & Xu, Y. (2020). Fabrication and characterization of cold-gelation whey protein-chitosan complex hydrogels for the controlled release of curcumin. Food Hydrocolloids, 103, 105619. 10.1016/j.foodhyd.2019.105619

11. Ma, J.-J., Yu, Y.-G., Yin, S.-W., Tang, C.-H., & Yang, X.-Q. (2018). Cellular Uptake and Intracellular Antioxidant Activity of Zein/Chitosan Nanoparticles Incorporated with Quercetin. Journal of Agricultural and Food Chemistry, 66(48), 12783–12793. 10.1021/acs.jafc.8b04571

12. Oakley, A. J. (2010). The structure of Aspergillus niger phytase PhyA in complex with a phytate mimetic. Biochemical and Biophysical Research Communications, 397(4), 745–749. 10.1016/j.bbrc.2010.06.024

13. Sandberg, A.-S., Hulthén, L. R., & Türk, M. (1996). Dietary Aspergillus niger Phytase Increases Iron Absorption in Humans. The Journal of Nutrition, 126(2), 476–480. 10.1093/jn/126.2.476

14. Saqib, Md. N., Khaled, B. M., Liu, F., & Zhong, F. (2022). Hydrogel beads for designing future foods: Structures, mechanisms, applications, and challenges. Food Hydrocolloids for Health, 2, 100073. 10.1016/j.fhfh.2022.100073

15. Steć, A., Chodkowska, M., Kasprzyk-Pochopień, J., Mielczarek, P., Piekoszewski, W., Lewczuk, B., Płoska, A., Kalinowski, L., Wielgomas, B., & Dziomba, S. (2023). Isolation of Citrus lemon extracellular vesicles: Development and process control using capillary electrophoresis. Food Chemistry, 424, 136333. 10.1016/j.foodchem.2023.136333

16. van den Berg, L., Toja Ortega, S., van Loosdrecht, M. C. M., & de Kreuk, M. K. (2022). Diffusion of soluble organic substrates in aerobic granular sludge: Effect of molecular weight. Water Research X, 16, 100148. 10.1016/j.wroa.2022.100148

17. Wang, S., Chen, K., Tian, A., Pan, M., Liu, X., Qu, L., Jin, J., Lv, S., Xu, Y., Li, Y., Yang, W., Zhang, X., Zheng, L., Zhang, Y., Yang, X., Zhong, F., Xu, L., & Ma, A. (2024). Effect of cooking methods on volatile compounds and texture properties in maize porridge. Food Chemistry: X, 22, 101515. 10.1016/j.fochx.2024.101515

18. Wong, S. K., Lawrencia, D., Supramaniam, J., Goh, B. H., Manickam, S., Wong, T. W., Pang, C. H., & Tang, S. Y. (2021). In vitro Digestion and Swelling Kinetics of Thymoquinone-Loaded Pickering Emulsions Incorporated in Alginate-Chitosan Hydrogel Beads. Frontiers in Nutrition, 8. https://www.frontiersin.org/articles/10.3389/fnut.2021.752207

19. Yan, J., Yin, L., Qu, Y., Yan, W., Zhang, M., Su, J., & Jia, X. (2022). Effect of calcium ions concentration on the properties and microstructures of doubly induced sorghum arabinoxylan/soy protein isolate mixed gels. Food Hydrocolloids, 133, 107997. 10.1016/j.foodhyd.2022.107997

20. Yang, E., Dong, H., Khongkomolsakul, W., Dadmohammadi, Y., & Abbaspourrad, A. (2024). Improving the thermal stability of phytase using core-shell hydrogel beads. Food Chemistry: X, 21, 101082. 10.1016/j.fochx.2023.101082

21. Yang, E., Yu, H., Choi, S., Park, K.-M., Jung, H.-S., & Chang, P.-S. (2021). Controlled rate slow freezing with lyoprotective agent to retain the integrity of lipid nanovesicles during lyophilization. Scientific Reports, 11(1), 24354. 10.1038/s41598-021-03841-4

22. Yang, J., Li, M., Wang, Y., Wu, H., Zhen, T., Xiong, L., & Sun, Q. (2019). Double Cross-Linked Chitosan Composite Films Developed with Oxidized Tannic Acid and Ferric Ions Exhibit High Strength and Excellent Water Resistance. Biomacromolecules, 20(2), 801–812. 10.1021/acs.biomac.8b01420

23. Zhang, Z., Zhang, R., Zou, L., & McClements, D. J. (2016). Protein encapsulation in alginate hydrogel beads: Effect of pH on microgel stability, protein retention and protein release. Food Hydrocolloids, 58, 308–315. 10.1016/j.foodhyd.2016.03.015

