## Supplementary material for "Core-Shell Hydrogel System to Protect the Enzyme Activity of Phytase from Environmental Stress": Tables, figures.

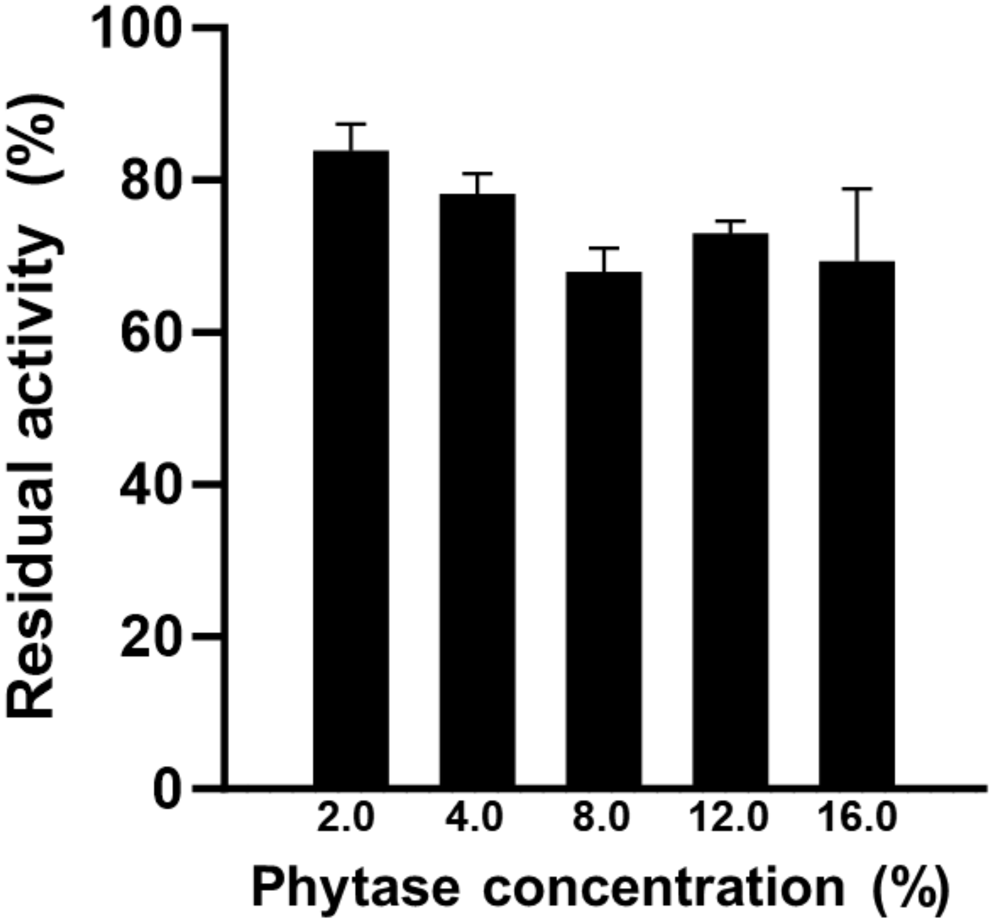


**Figure S1.** Thermal stability of phytase A-loaded core-shell hydrogel beads, depending on the phytase concentration.

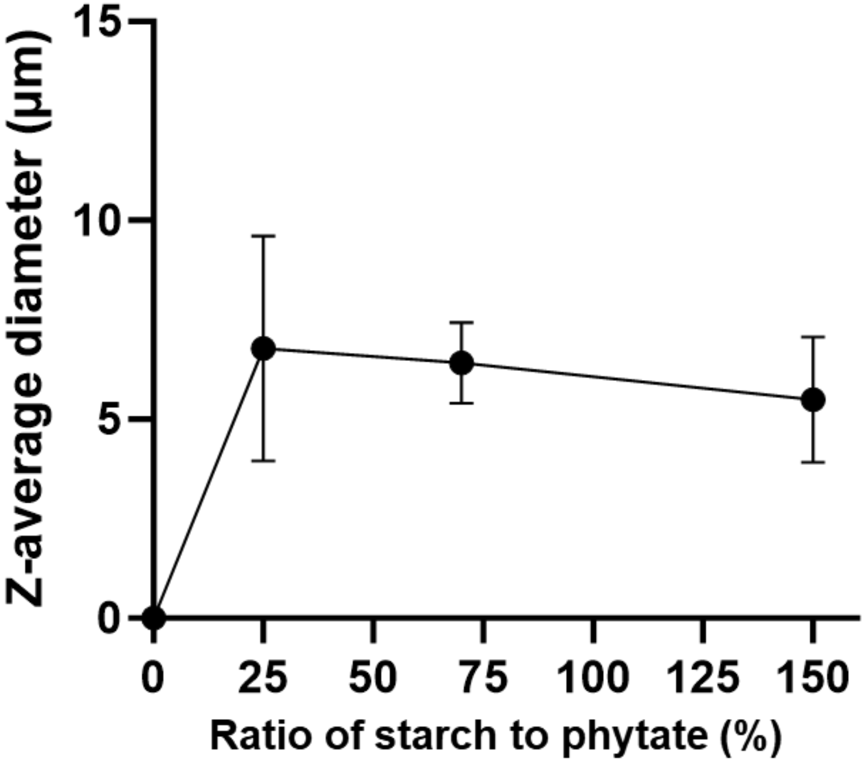


**Figure S2.** Z-average diameter of phytate-starch complex depending on the ratio of starch to phytate.
